# Human genetic variants associated with COVID-19 severity are enriched in immune and epithelium regulatory networks

**DOI:** 10.1101/2021.12.17.473140

**Authors:** Zhanying Feng, Xianwen Ren, Zhana Duren, Yong Wang

## Abstract

Human genetic variants can influence the severity of symptoms being infected with SARS-COV-2. Several genome-wide association studies have identified human genomic risk SNPs associated with COVID-19 severity. However, the causal tissues or cell types of COVID-19 severity are uncertain and candidate genes associated with these human risk SNPs were investigated in genomic proximity instead of their functional cellular contexts. Here, we compiled regulatory networks of 77 human contexts and revealed those risk SNPs’ enriched cellular contexts and associated transcript factors, regulatory elements, and target genes. Twenty-one human contexts were identified and grouped into two categories: immune cells and epithelium cells. We further aggregated the regulatory networks of immune cells, epithelium cells, and immune-epithelium crosstalk and investigated their association with risk SNPs’ regulation. Two genomic clusters, chemokine receptors cluster and OAS cluster, showed the strongest association with COVID-19 severity and different regulations in immune and epithelium contexts. Our findings were supported by analysis on both microarray and whole genome sequencing based GWAS summary statistics.

## Introduction

Coronavirus disease 2019 (COVID-19) is an infectious disease caused by the SARS-CoV-2 virus. Most people infected with the virus will experience mild to moderate respiratory illness and recover without requiring special treatment. However, some people will become seriously ill and require medical attention. Scientists from all over the world have made great efforts to understand the genetic mechanism of COVID-19 severity, which may lead to efficient prevention stratagem and effective cure. Genome-wide association analysis (GWAS) has been the ideal option to find human genetic variants associated with severity phenotypes. Recently, applying GWAS to phenotypes of COVID-19, such as infection and severe respiratory symptoms, has helped us identify some risk loci associated with COVID-19. For example, the “The Severe COVID-19 GWAS Group” conducted a GWAS involving 1,980 patients with severe COVID-19 symptoms and detected SNP rs11385942 at locus 3p21.31 and rs657152 at locus 9q34.2^1^. Erola P. C. *et al*. used GWAS to study 2,244 critically ill COVID-19 patients from 208 UK intensive care units (ICUs) and found rs10735079 on 12q24.13 in a gene cluster encoding antiviral restriction enzyme activators (*OAS1, OAS2, OAS3*), rs2109069 on 19p13.2 near the gene *TYK2*, rs2109069 on 19p13.3 within *DPP9*, and rs2236757 on 21q22.1 in the interferon receptor gene *IFNAR2*^2^. “The COVID-19 Host Genetics Initiative” conducted several GWAS, including “very severe respiratory confirmed COVID” and “hospitalized COVID”, and also found many potential loci^3^. These identified human risk loci allowed us to further understand the underlying mechanism of COVID-19 severity.

Interpreting these COVID-19 associated human genetic variants remains challenging since most genetic variants were located in the non-coding regulatory region with high linkage disequilibrium^4-6^ and were functional in time- and space-specific contexts^7^. This implies that we must elucidate these genetic variants in the proper cellular contexts. Several works have been done to uncover the regulatory mechanism of genetic variants. For example, FUMA^8^ and GREAT^9^ used the routine way to link SNPs to the nearby genes. SMR^10^ and Sherlock^11^ utilized the expression information to find causal SNP affecting gene regulation. However, these methods didn’t consider either the direct cis-regulation or the regulatory effects in biological network. Recently, we developed a method, PECA, to infer context specific regulatory network with paired expression and chromatin accessibility data from diverse cellular context^12^ and then extended to PECA2 to reconstruct regulatory network for a single sample^13^. The reconstructed regulatory networks have been successfully applied to reveal critical regulations for time course data^14^. We noticed that the availability of the public available paired expression and chromatin accessibility data, such as ENCODE and ROADMAP, ensured the diversity in tissue source of the cells and the germ layer lineage representation. This allows us to run PECA2 to systematically construct a regulatory network atlas of 77 human contexts, which served as a valuable resource for genetic variants interpretation in multi-cellular contexts.

Here, we utilized these 77 regulatory networks to interpret the SNPs associated with COVID-19 severity. We found the relevant tissues of COVID-19 severity were categorized into two main cell types: immune cells and epithelium cells. Then in these two cell type categories, we illustrated the detailed SNP associated regulations: SNP located in regulatory elements (REs), their upstream transcription factors (TFs), and their downstream target genes (TGs). We found that two gene clusters (chemokine receptors and OAS cluster) showed different regulation patterns in two cell type categories. Our COVID-19 severity associated regulatory network will be promising to serve as a valuable perspective to study COVID-19.

## Results

### 1. Human genetic variants of COVID-19 severity are enriched in immune and epithelium cells

We first collected paired expression and chromatin accessibility data from 76 human contexts and constructed a human regulatory network atlas. The samples covered all three germ layers, such as frontal cortex (ectoderm), primary T cells (mesoderm), and upper lobe of left lung (endoderm). With each sample’s paired expression and chromatin accessibility data as input, we used PECA2 model^13^ to construct regulatory network for each context (**Methods**). The basic unit of regulatory network is TF-REs-TG triplet and each triplet denotes that a TF binds on the REs and regulates a nearly TG. We also included the regulatory network of cranial neural crest cells^15^ (CNCC), a migratory cell population in early human craniofacial development, into our regulatory network atlas. Then we applied these regulatory networks of 77 human contexts to interpret genetic variants of COVID-19 severity.

We fetched the 542 SNPs with significant associations (P≤ 5 × 10^−8^) with “very severe respiratory confirmed COVID” (“A2_ALL” study in “The COVID-19 Host Genetics Initiatives”). To evaluate the relevance between SNPs and 77 human contexts, the fold enrichment (FE) score of these 542 SNPs in RE sets of 77 human contexts were calculated (**Methods**). We set the threshold of FE score 3.0 and found that 21 contexts were relevant to COVID-19 severity, such as “primary monocytes” and “upper lobe of left lung” (**Figure 1A**). Furthermore, these 21 contexts could be classified into two categories. The first category was immune cells, including 10 cell types: “Primary monocytes”, “Primary B cells”, “Jurkat”, “GM12878”, “Primary natural killer cells”, “CD4 primary cells”, “Fetal thymus”, “Primary T cells”, “CD8 primary Cells”, and “Hematopoietic multipotent progenitor cell”. The second category was epithelium cells and consisted of 11 cell types: “Vagina”, “Omental fat pad”, “Gastric”, “Upper lobe of left lung”, “Lower leg skin”, “Esophagus squamous epithelium”, “Esophagus muscularis mucosa”, “Fetal spleen”, “Subcutaneous adipose tissue”, “Spleen”, and “T47D”.

**Figure 1.**
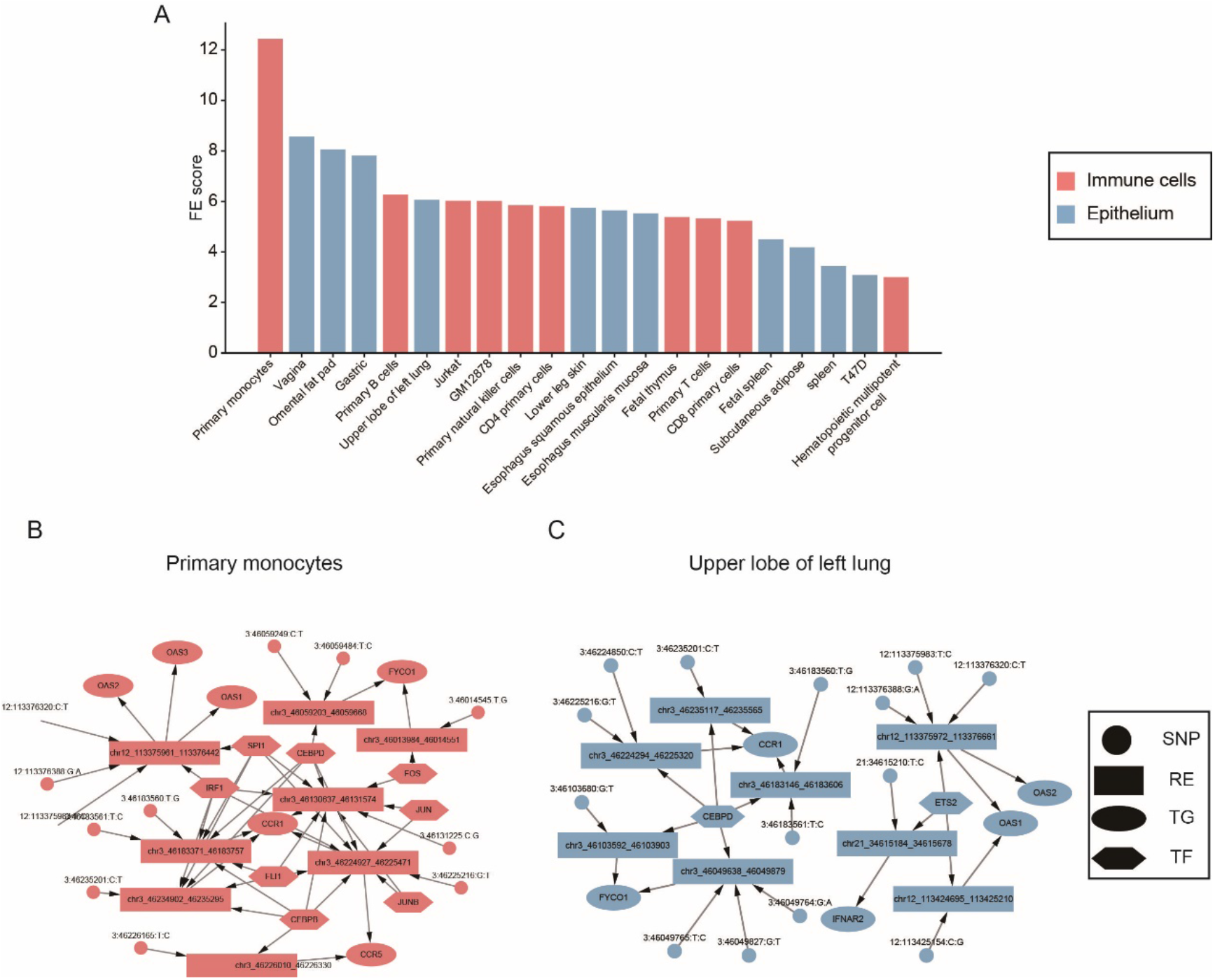
(A). 21 COVID-19 severity relevant contexts ranked by fold enrichment (FE) scores. Red: immune cells. Blue: epithelium cells. (B). COVID-19 SNP associated regulatory of “Primary monocytes”, which is an immune cell type. (C). COVID-19 SNP associated regulatory of “Upper lobe of left lung”, which is an epithelium cell type.

We further extracted sub-network that was associated with SNPs of COVID-19 severity in every relevant context (**Methods**). For example, “Primary monocytes” was one of the immune cell types. And 12 SNPs were located in the 8 REs in regulatory network of “Primary monocytes” (**Figure 1B**). These REs were predicted to regulate 9 TGs, such as *CCR1* and *FYCO1. CCR1* played a key role in T-cell-mediated respiratory inflammation^16^. These REs were bound by immunity associated TFs, such as *CEBPB*/*D*^17-20^ and *FLI1*^21^. On the other hand, “Upper lobe of left lung” was one of the epithelium cell type (**Figure 1C**), and 8 REs were associated with 14 SNPs of COVID-19 severity and regulated 5 TGs, such as *CCR1* and *OAS1*. Polymorphisms of *OAS1* have been reported to affect susceptibility to a variety of viral diseases^22,23^. Some TFs regulating these REs were linked to lung epithelium, such as *CEBPD*^24^ and *ETS2*^25^.

In summary, we built a regulatory network atlas of 77 human contexts and leveraged their regulatory elements to reveal relevant contexts to human genetic variants of COVID-19 severity. Two categories of cell types were found: the first was epithelium cells and may be involved with susceptibility to viral diseases; the other category was immune cells and possibly linked to severity after being infected with disease. The relevance to two categories of cell types were reproduced by more recent GWAS with a larger cohort from HGI (**Figure S1**).

### 2. Construction of SNP associated regulatory network of immune and epithelium cells

To further understand SNPs’ function in two categories of cell types, we constructed COVID-19 severity associated regulatory networks in immune cells and epithelium cells respectively. Briefly, we pooled the SNPs associated networks of 10 immune cell types into a COVID-19 severity associated regulatory network of immune cells. Similarly, we pooled the SNPs associated regulatory network of 11 epithelium cell types into a COVID-19 severity associated regulatory network of epithelium cells (**Methods**).

In COVID-19 severity associated immune regulatory network, there were totally 17 TFs, 25 REs and 15 TGs associated with 38 SNPs (**Figure 2A**). This sub-regulatory network was highly associated with immune functions. For example, *TCF3/7/12* were important upstream TFs in regulatory network of immune cells and played vital roles during T-cell development^26,27^. *IRF1/3/4* were critical TFs in the cellular differentiation of hematopoietic cells and in the regulation of gene expression in response to pathogen-derived danger signals^28^. *CCR5* was downstream TGs in this regulatory network, and it encoded a protein on the surface of white blood cells that is involved in the immune system^29^. We found that 2 TGs (*CCR5, CXCR6*) of the 15 immune TGs were also differentially expressed in blood samples of fatal COVID-19 patients^30^ (*P* ≤ 0.014), which supported that our reconstructed sub-regulatory networks of immune cells were significantly associated with severity of COVID-19. In COVID-19 severity associated epithelium regulatory network, there were totally 16 TFs, 24 REs and 10 TGs associated with 42 SNPs (**Figure 2B**). This SNP associated regulatory network was involved with the functions of epithelium cells. For example, *ETS2*, which was core TF in regulatory network of epithelium cells, could promote epithelial-to-mesenchymal transition in renal fibrosis^25^. *ELF3* was also essential for mesenchymal to epithelial transition^31^. *NBEAL2* was a TG in epithelium regulatory network and the *Nbeal2*-deficient mice exhibited impaired development of functional granulation tissue due to severely reduced differentiation of myofibroblasts in the model of excisional skin wound repair^32^. These literature evidence showed that our reconstructed SNP associated regulatory networks were promising to reveal COVID-19 severity associated regulations in immune and epithelium cells.

**Figure 2.**
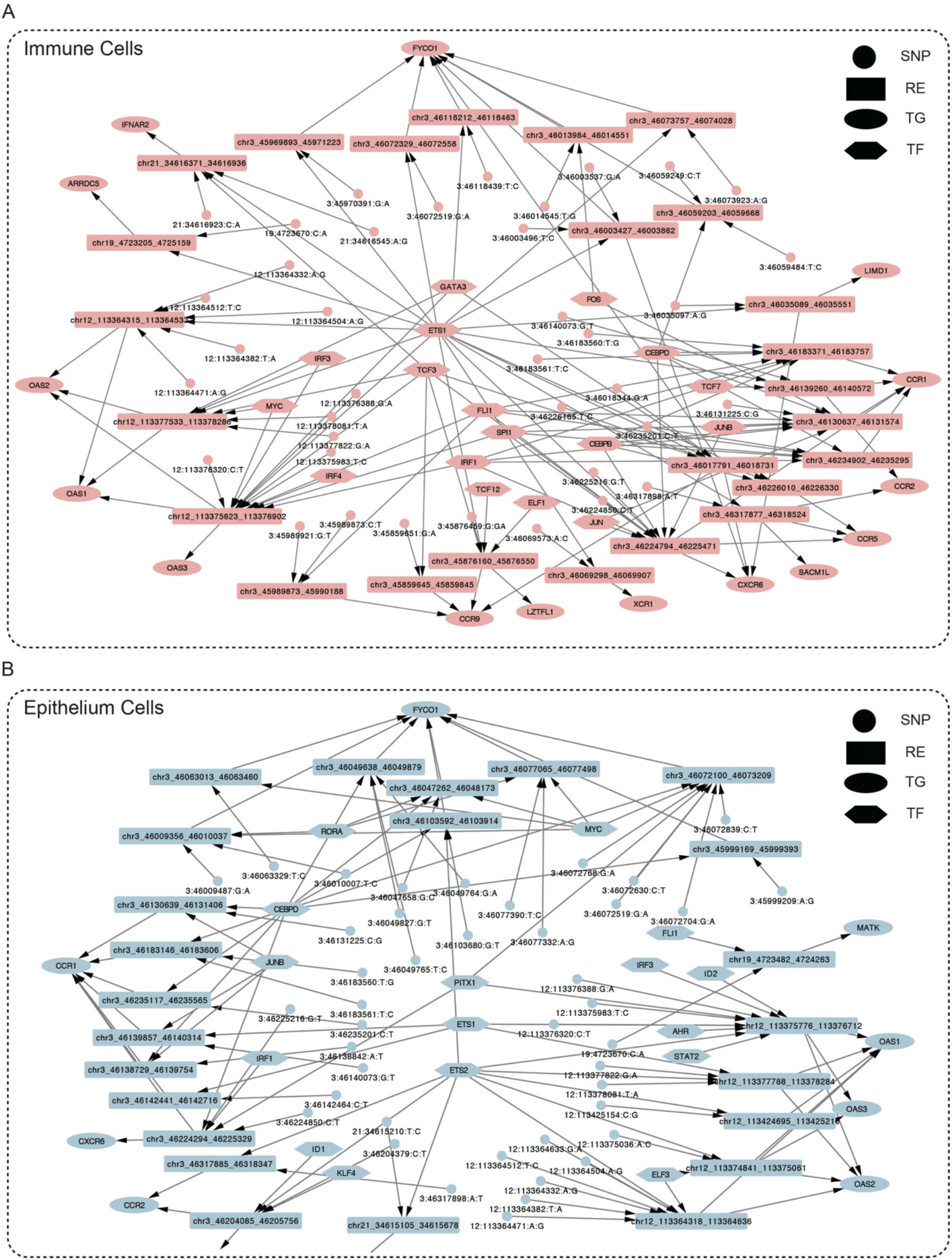
(A). COVID-19 severity associated regulatory network of immune cells. (B). COVID-19 severity associated regulatory network of epithelium cells.

We further overlapped our SNP associated regulatory network with eQTL dataset. Our SNP associated regulatory network revealed the link between SNPs and genes by SNP’s location in REs. For example, SNP “12:113364332:A:G” was located in RE “chr12:113364318-113364636” and “chr12:113364318-113364636” regulated *OAS1*, which gave the association between SNP “12:113364332:A:G” and gene *OSA1*. In this way, we totally obtained 91 SNP-TG association from immune and epithelium regulatory networks. Then we collected significant variants-gene association of 49 tissues from GTEx v8. We found that our SNP-TG associations were highly reproducible by eQTL variants-gene association. First, there were 78 (85.7%) SNP-TG associations could be validated as an eQTL variant-gene association in at least one tissue. On the other hand, when we ranked the 49 tissues by the overlapping number of eQTL with our SNP-TG association, we found that the top ranked tissues were also immune and epithelium cell types (**Figure 3**). For example, “esophagus mucosa”, which was one of epithelium cell types, ranked first among 49 tissues. There were some other tissues that were related to epithelium cells, such as “Cells Cultured fibroblasts”, “Skin Not Sun Exposed Suprapubic”, “Spleen”, “Skin Sun Exposed Lower leg”, “Lung”. And there were also immune tissues among the top ranked tissue, such as “Artery Tibial” and “Whole Blood”. To check if our SNP-TG associations were causally responsible for COVID-19 severity, we conducted colocalization analysis with GWAS summary statistics of COVID-19 severity and eQTL form GTEx by SMR^10^. We obtained 57 SNP-TG pairs in 49 tissues and found that 2 of our 91 SNP-TG association were validated by the colocalization analysis, which was significantly overlapped (*P* = 1 × 10^−7^). These two SNP-TG association were related to *OAS3*, revealing that “12:113376320:C:T” and “12:113375983:T:C” might causally regulate expression of *OAS3* and influenced severity of COVID-19 symptoms. These results showed that our COVID-19 severity associated regulatory network could be supported by independent eQTL dataset.

**Figure 3.**
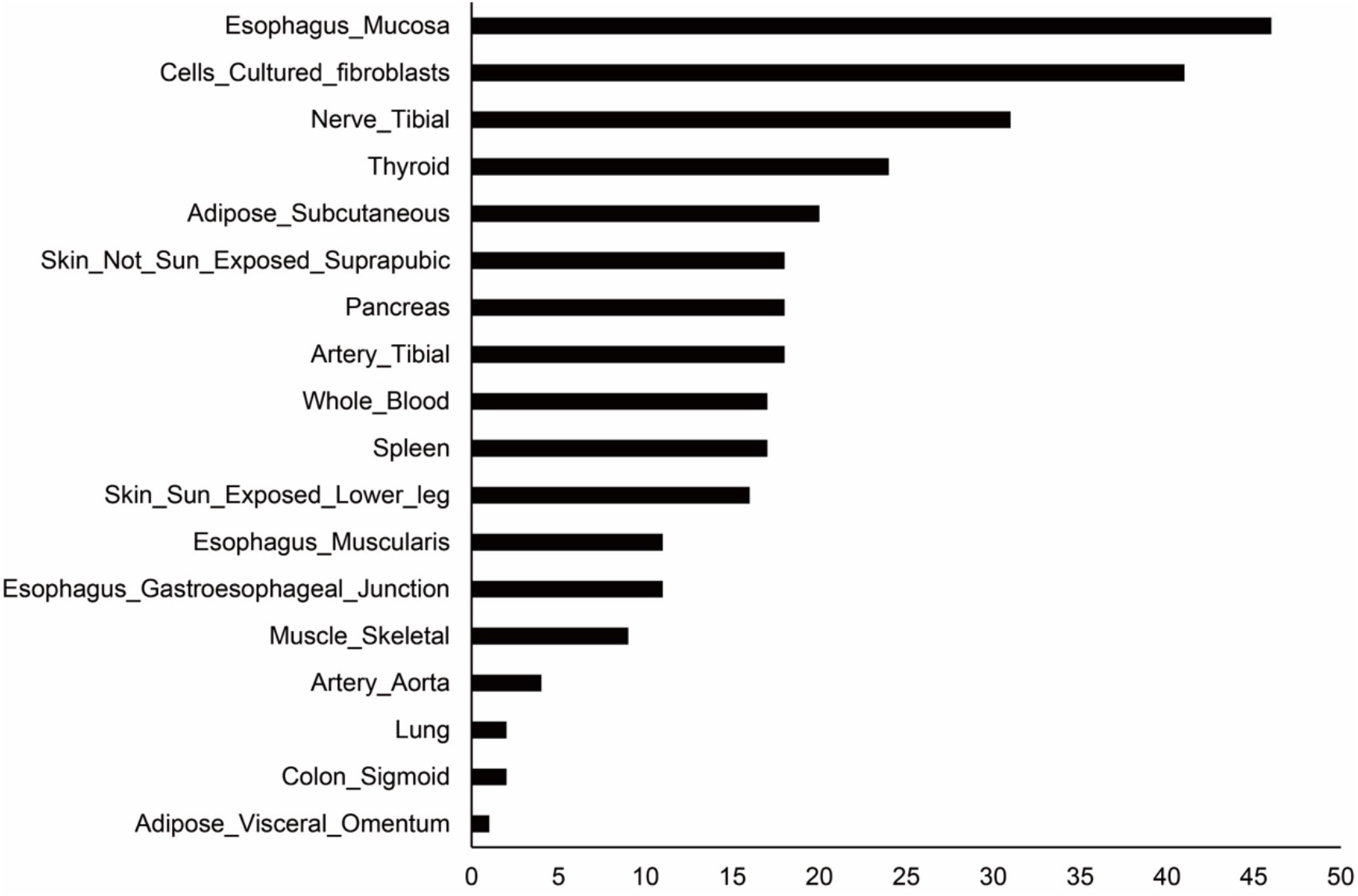
Overlapping of SNP-TGs inferred from our regulatory networks with eQTL of 49 tissues in GTEx. The tissues are ranked by the number of eQTL overlaps.

In summary, we have constructed COVID-19 severity associated regulatory network of immune cells and epithelium cells. These two regulatory networks were tightly linked to immune and epithelium functions and could be validated by eQTL of immune and epithelium tissues in GTEx.

### 3. Associating two cell types’ regulatory networks revealed shared regulatory structure

After reconstructing and validating the regulatory networks of immune and epithelium cells, we next compared these two cell types to find conservation and divergence. We first compared the TFs, TGs, REs, and SNPs in the regulatory networks of immune and epithelium cells. For TFs, 7 TFs were shared by two cell types, such as *ETS1* and *IRF1*. There were 10 immune-specific TFs (such as *TCF7* and *ELF1*) and 9 epithelium-specific TFs (such as *ETS2* and *KLF4*). There were also 8 overlapped TGs (such as *CCR1, FYCO1*), 7 immune-specific TGs (such as *CCR5* and *LZTFL1*) and 2 epithelium-specific TGs (*MATK* and *NBEAL2*). For SNPs and REs, about half of them in immune regulatory network were shared by epithelium regulatory network (**Figure 4**).

**Figure 4.**
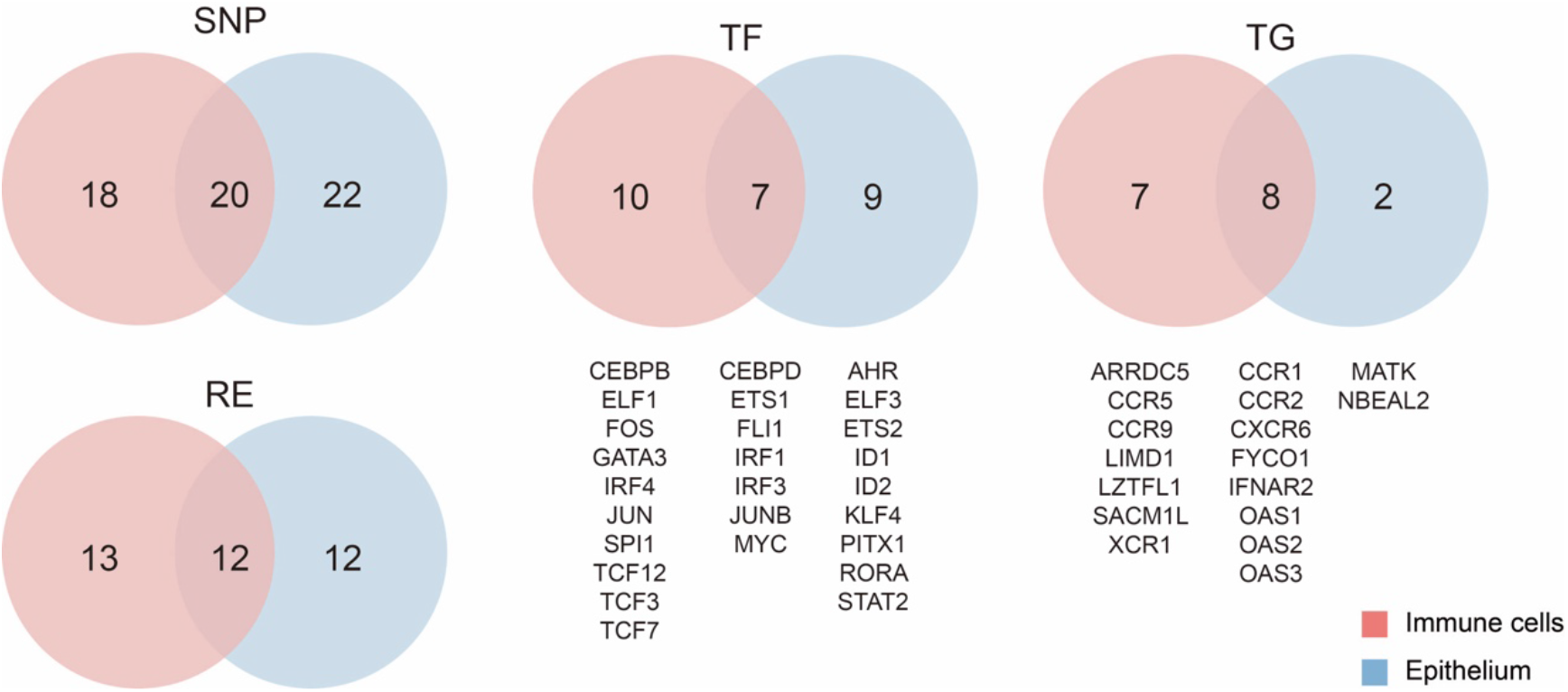
Overlapping of SNP, RE, TF, TG between immune regulatory network and epithelium regulatory network.

We found that the regulatory network could be clustered into four clusters according to their location and this clustering was shared between immune and epithelium cells (**Figure 2A, B**). The first cluster was on chromosome 3. This cluster was involved with regulation of *CCR1/2/5/9, CXCR6, FYCO1, LIMD1, LZTFL1, SACM1L, XCR1* in immune cells and *CCR1/2/5/9, CXCR6, NBEAL2* in epithelium cells. The second cluster was located in chromosome 12 and involved with regulation of *OAS1/2/3* in both immune and epithelium cells. The third cluster was in chromosome 19. In immune cells, this cluster was associated with *ARRDC5* and in epithelium cells, it was associated with *MATK*. The last cluster was in chromosome 21 and linked to *IFNAR2* in two cell types. These four genetic variants associated regulatory clusters may exert genetic influence on infection and severity of COVID-19.

In the regulatory network of two cell types, we found two classes of upstream TFs (**Figure 2A, B**). The first class of TFs regulated most of the four regulatory clusters. For example, in regulatory network of immune cells, *ETS1, TCF3* and *GATA3* were involved with at least three regulatory clusters. And for the epithelium cells, *ETS1, ETS2*, and *PITX1* were classified into this class. Contrary to the broad regulation of the first class, the second class of TFs was only responsible for a small part of regulatory network. For example, in immune regulatory network, *CEBPD, CEBPB, TCF7, FOS, JUNB, JUN, ELF1* were only regulate REs in chromosome 3 regulatory cluster. And *IRF3, IRF4*, and *MYC* only responsible for regulations in chromosome 12 regulatory cluster. In the epithelium regulatory network, *CEBPD, JUNB, IRF1, ID1, RORA, MYC*, and *KLF4* were only involved with the regulatory of chromosome 3 regulatory cluster. *ID2, IRF3, AHR, STAT2, ELF3* was associated with the chromosome 12 regulatory cluster.

Through the comparison of two cell types’ regulatory networks, we found these two cell types shared many TFs, TGs, REs, and SNPs. The regulatory structure (four regulatory clusters based on location and two classes of upstream TFs) were conserved in two cell types. Despite the shared regulatory structure, there were also many cell-type-specific TFs, TGs, REs, and SNPs.

### 4. Chromosome 3 and 12 clusters showed distinct regulatory program in immune and epithelium cells

From the above analysis of regulatory structure, we noticed that there are two relatively denser clusters in immune and epithelium regulatory: chromosome 3 and chromosome 12 clusters, indicating that they may play an essential role in COVID-19 severity. Here we focused on these two regulatory clusters.

For regulatory cluster in chromosome 12, these regulatory programs were mainly involved with *OAS* cluster (*OAS1/2/3*) in ten immune cell types and nine epithelium cell types. These *OAS* cluster was closely associated with immune and epithelium functions. For example, *OAS1* polymorphisms played potential roles in respiratory infection from human bronchial epithelial cells^23^. The *OAS2* protein is a well-known innate immune activated antiviral enzyme catalyzing synthesis of 2’-5’-oligoadenylate for RNase L activation and inhibition of viral propagation^33^. We found that the REs of this cluster in different cell types were mainly distributed around the promoter of *OAS3* (**Figure 5A**) and there were many COVID-19 severity associated SNPs within these REs, which was consistent among immune and epithelium cell types. To add additional evidence of regulation in this area, we collected previously published promoter-capture Hi-C (PCHi-C) data of primary blood cell types^34^. We used the PCHi-C interactions that have CHiCAGO scores ≥ 5 in at least one analyzed cell type and found there was a loop between promoter of *OAS3* and promoter of *OAS2*. This observation induced a hypothesis that the REs around the promoter of *OAS3* were cis-regulatory elements of *OAS2* in immune cells. To validate this hypothesis, we first checked the REs regulating the three OAS genes in 10 immune cell types and found that all 15 REs were predicted to regulate *OAS2*, 13 REs were predicted to regulate *OAS1* and only 7 REs were predicted to regulate *OAS3*, which revealed that more evidence supported that REs in this locus regulated *OAS2*. Then we checked the expression of three *OAS* genes and found *OAS2*’s expression was the highest among *OAS* cluster in immune cell types (**Figure 5B**). We also computed the averaged correlation between the openness score of REs in *OAS3*’s promoter and expression of three OAS genes across 148 samples of human (**Table S1**). The result showed that the accessibility of these REs was more correlated to expression of *OAS2* (**Figure 5C**). In summary, these evidence of PCHi-C loop, regulatory network, expression data, and RE-TG correlation indicated that in immune cells, the REs around *OAS3*’s promoter were more likely to regulate *OAS2*.

**Figure 5.**
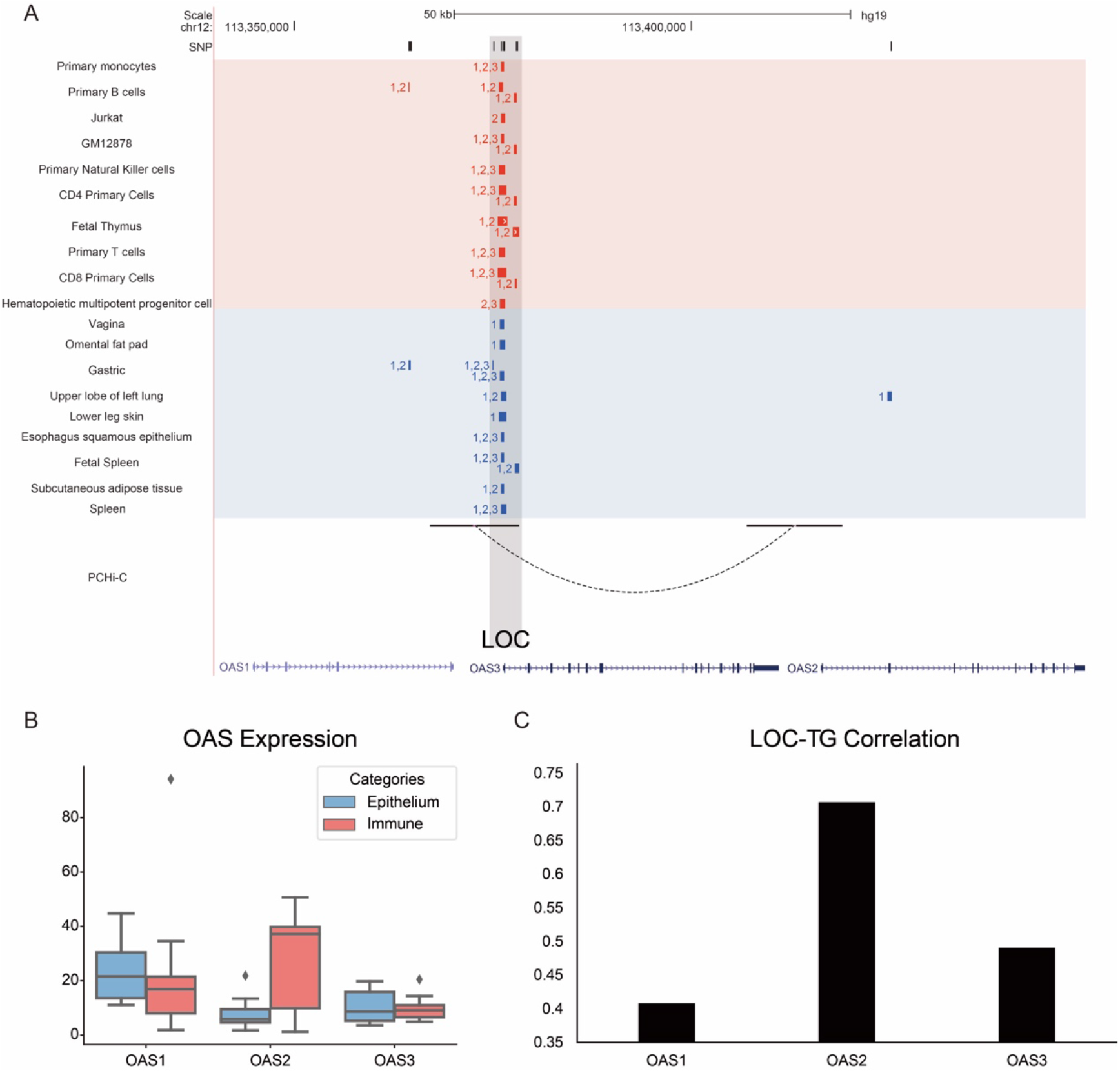
(A). SNPs, REs and PCHi-C loop associated with *OAS1/2/3*. The RE tracks consisted of 11 immune cell types and 9 epithelium cell types. PCHi-C loops were from blood samples. The number labels around the RE were indicator of RE’s target genes. (B). Expression of *OAS1/2/3* in immune and epithelium cells. (C). Correlations between accessibility of REs and expression of *OAS1/2/3*.

For regulatory cluster in chromosome 3, these regulatory programs were mainly involved with chemokine receptor (*CCR1/2/5/9, CXCR6*, and *XCR1*) in eight immune cell types and eight epithelium cell types. In literature, there were many reports that these chemokine receptors were associated with the function of both immune and epithelium cells^16,29^. We found five main regulatory areas that were associated with SNPs and regulated these chemokine receptors (we name them as LOC1-LOC5, **Figure 6A**). Some of these REs’ regulation was shared by two cell types. For example, LOC3 were predicted to regulate *CCR1* in both immune and epithelium cells. And there was also a PCHi-C loop between LOC3 and *CCR1*. The other LOCs’ regulatory programs were different. For example, LOC1 and LOC2 were immune-specific REs and regulated *CCR9* and *CXCR6*. While REs in LOC4 were shared by two cell types, they were predicted to regulate *CCR5* in immune cells but regulate *CCR1* in epithelium cells. LOC4 and *CCR5* were also contacted by a PCHi-C loop in blood cells. Similar to LOC4, LOC5 regulated *CCR2/5* in immune cells but only regulate *CCR2* in epithelium. The expression of these chemokine receptors supported the regulation above: *CCR1/2/5/9* and *CXCR6* were relatively high expressed in immune cells but only *CCR1* and *CCR2* were expressed in epithelium cells (**Figure 6B**). We also checked the RE-TG correlation and found the above regulations were also supported. LOC1 showed the highest correlation with *CCR9*. LOC2 were more correlated with *CXCR6* and *CCR9*. LOC3 was most associated with *CCR1*. LOC4 showed higher correlation with *CCR1* and *CCR5*. And LOC5 was correlated with *CCR5* (**Figure 6C**).

**Figure 6.**
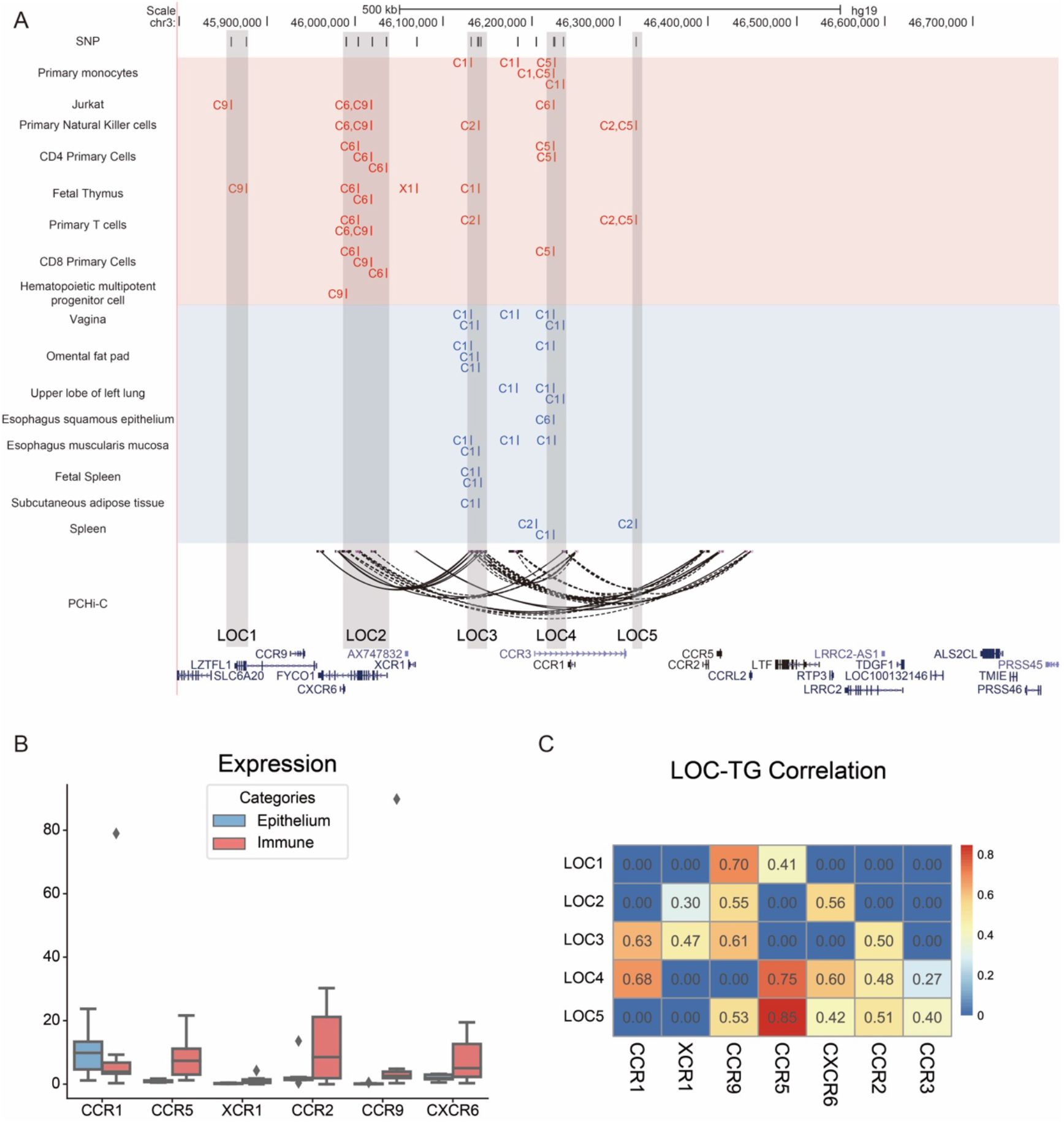
(A). SNPs, REs and PCHi-C loop associated with chemokine receptors. The RE tracks consisted of 11 immune cell types and 9 epithelium cell types. PCHi-C loops were from blood samples. The labels around REs were indicator of TGs: C1-CCR1, C2-CCR2, C5-CCR5, C6-CXCR6, C9-CCR9, X1-XCR1. (B). Expression of chemokine receptors in immune and epithelium cells. (C). Correlations between accessibility of REs and expression of chemokine receptors.

Taken together, we found some shared and distinct regulatory programs in immune and epithelium by detailed study of two main COVID-19 severity associated regulatory cluster on chromosome 3 and 12. These conserved and divergent regulations may give biological insights about how virus infected human through epithelium cells and how human defensed SARS-COV-2 with our immune system.

### 5. Analyzing whole genome sequencing based GWAS summary statistics reproduced relevant immune and epithelium cells and refined the regulatory networks

Microarray genotyping based genome-wide association studies (GWAS) have been the most common approach for identifying disease associations across the whole genome and can interrogate over 4 million markers per sample. Whole-genome sequencing provides a comprehensive base-by-base method for interrogating the 3.2 billion bases of the human genome. Higher resolution GWAS summary statistics hold the promise to reveal richer and more accurate genetic variants. We noticed that Kousathanas *et al*. used whole genome sequencing data (WGS) of a larger cohort to conduct GWAS analysis and found 1,322 SNPs (P≤ 5 × 10^−8^) with significant association with severity with COVID-19^35^. We then used this independent data to validate and refine our relevant contexts and regulatory networks.

We first conducted the SNP enrichment analysis on these 1,322 significant SNPs to find relevant contexts. Eighteen cellular contexts obtained FE score ⩾3.0 and were grouped into two categories: immune cells such as “Primary monocytes”, “CD4 primary cells” and epithelium cells such as “Upper lobe of left lung”. This results faithfully reproduced our results in microarray genotyping based GWAS data that human genetic variants of COVID-19 severity were enriched in immune and epithelium cells. Furthermore, this WGS genotyping based GWAS also revealed that immune cells were more associated with severity of COVID-19 that epithelium cells (**Figure 7A**). Next, we sought to refine our regulatory networks of immune and epithelium cells with the overlapped SNPs of the WGS based GWAS and previous microarray based GWAS. We found that among the 38 SNPs of the previous reconstructed regulatory network of immune cells, 25 SNPs were reproducible in the WGS based GWAS. These 25 SNPs formed a regulatory network of immune cell, which include 12 TGs such as *CCR1/2/5/9*, 14 TFs such as *FLI1, JUNB, CEBPD*, and 20 REs (**Figure 7B**). And for epithelium cells, we found that among the 42 SNPs of the previous reconstructed regulatory network, 22 SNPs were also significant in the WGS based GWAS. These 22 SNPs gave a regulatory network of epithelium cell, which include 6 TGs such as *CCR1/2, FYCO1*, 14 TFs such as *ETS1/2, KLF4*, and 16 REs (**Figure 7C**).

**Figure 7.**
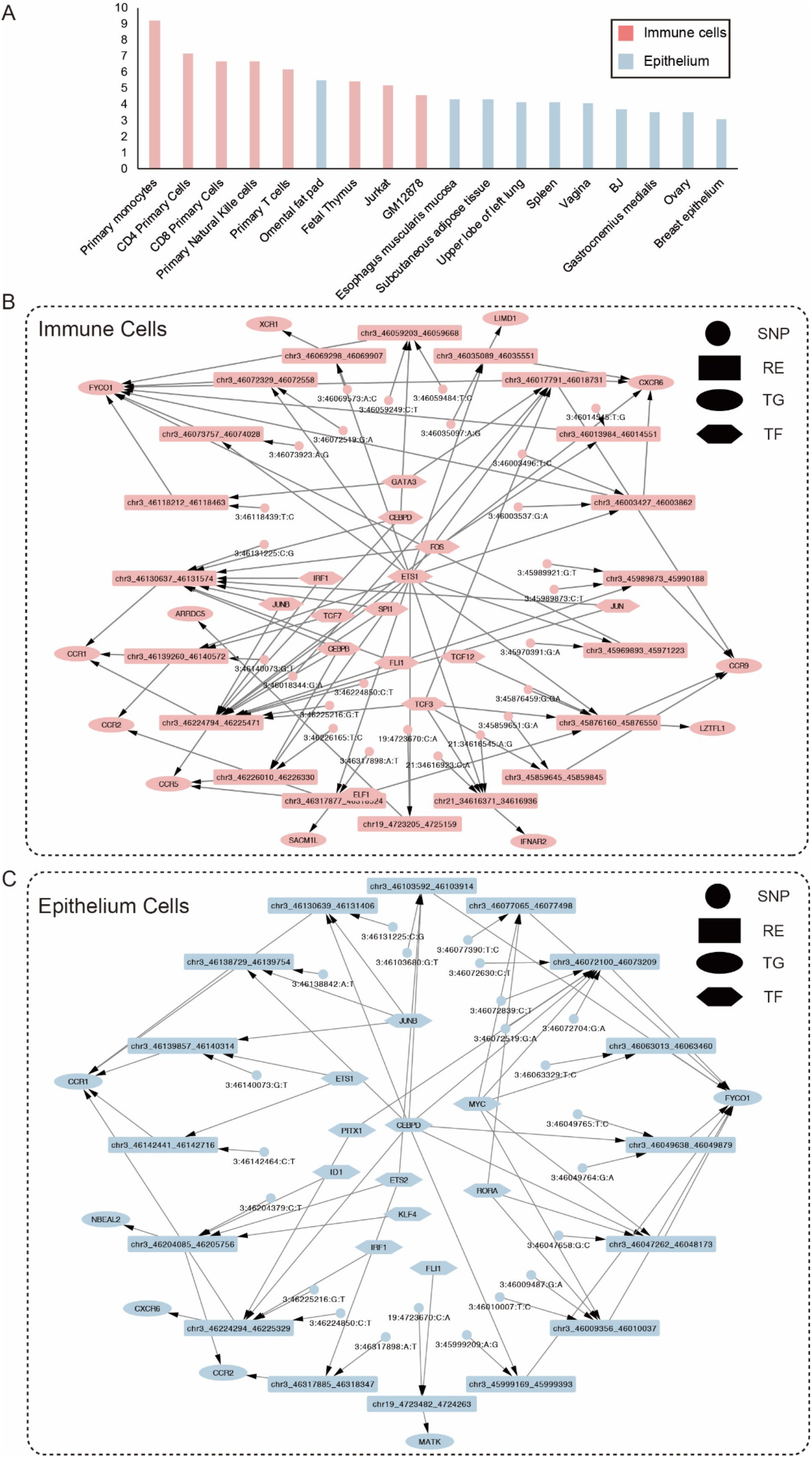
(A). Analyzing whole genome sequencing based GWAS summary statistics revealed 18 COVID-19 severity relevant contexts, which are ranked by fold enrichment (FE) scores. Red: immune cells. Blue: epithelium cells. (B). Refined COVID-19 severity associated immune regulatory network. (C). Refined COVID-19 severity associated epithelium regulatory network.

In summary, we conducted regulatory network analysis with a recent published WGS based GWAS and validated our findings that human genetic variants of COVID-19 severity were enriched in immune and epithelium cells. We also used the reproducible SNPs to obtain refined regulatory networks of immune and epithelium cells.

## Discussion

The COVID-19 pandemic has influenced human’s life all over the world. Scientists made their efforts to gain more understanding of pathology of COVID-19 and to find effective cure. GWAS is a powerful tool to discover suspicious loci for interesting traits. By profiling phenotype and genotype of COVID-19 infected people, the “COVID-19 host genetics initiative” had conducted GWAS analysis with more and more power^36^. Through these GWAS, many risk loci that may be associated with COVID-19 infection and severity were found. Following these discoveries, we used regulatory network atlas to interpret these risk loci and sought to gain biological insights of COVID-19.

First, we found that COVID-19 severity associated SNPs were mainly enriched in two types of human contexts: immune cells and epithelium cells. We used another GWAS of COVID-19 severity with larger cohort (https://www.covid19hg.org/results/r6/) to show that the association with immune cells and epithelium cells were quite reproducible (**Figure S1**). We then reconstructed the COVID-19 severity associated regulatory network in the two major contexts. The COVID-19 severity associated regulatory networks were validated by recently published RNA-seq data of COVID-19 patients and eQTL from GTEx. And further analysis revealed regulatory structure of COVID-19 associated regulatory networks: the four regulatory clusters and two classes of upstream TFs. The regulatory structure showed both conservation and divergence between two of cell types. Then we focused on two important regulatory clusters (OAS cluster and chemokine receptor cluster) and revealed the causal genes and differential regulation between two cell types. Finally, analysis on a recent WGS based GWAS with a larger cohort validated the two COVID-19 severity’s relevant cell types and refined the associated regulatory networks.

Several factors hindered the post-GWAS analysis of COVID-19. First of all, one more powerful GWAS with larger cohort is in need for such a complex trait. In this paper, the GWAS only detected 542 SNPs that significantly associated with COVID-19’s symptoms, which is not comparative to normal phenotype, such as height or BMI. Some efforts have been made to improve the power of GWAS. For example, the most recent GWAS conducted by “COVID-19 host genetics initiative” has included more than 8,000 very severe respiratory confirmed cases into analysis. On the other hand, a more comprehensive regulatory network atlas is promising for better understanding of COVID-19 severity. Although our current regulatory network atlas has covered many tissues, it is still far from complete and is limited in cell type resolution. As the symptoms of COVID-19 are involved with many organs, a more comprehensive regulatory network atlas, even at cell type level from single cell multi-omics data, will interpret more genetic variants associated with COVID-19.

## Methods

### Construction of regulatory network from paired expression and chromatin accessibility data

We utilized PECA2 to infer genome-wide and context-specific regulatory networks based on gene expression and chromatin accessibility data in that context^13^. Given paired RNA-seq and ATAC-seq data for a sample, PECA2 hypothesized that TF regulated the downstream TG by binding at REs. The regulatory strength of a transcription factor (TF) on a target gene (TG) was quantified by the trans-regulation score, which was calculated by integrating information from multiple regulatory elements (REs) that may mediate the activity of the TF to regulate the TG. A prior TF-TG correlation across external public data from ENCODE database was included in the trans-regulation score definition to distinguish the TFs sharing the same binding motif (i.e., TFs from the same family).

We collected paired expression and chromatin accessibility data of 76 human tissues or cell lines (**Table S2**) and applied PECA2 to these data to obtain regulatory networks of 76 human contexts. Furthermore, we recently constructed high-quality regulatory network of cranial neural crest^15^ and we included it into our analysis to form regulatory network atlas of 77 human contexts.

### Fold enrichment score of SNPs in given region set

Given a group of SNPs and a RE set, we defined the fold enrichment (FE) score as follows,

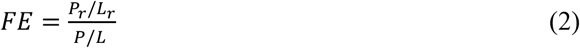

Where *P*_*r*_ was the number of SNPs in REs. *L*_*r*_ was the length of the REs. *P* was the total number of SNPs. *L* was the genome length.

We calculated the *FE* score of 542 SNPs of COVID-19 severity in RE sets of 77 human contexts and used a criterion of *FE* ≥ 3 to find 21 contexts that associated with COVID-19 severity. The FE score was also computed in the 1,322 SNPs of a recent GWAS. Under the same threshold, 18 contexts were found to be relevant to severity of COVID-19.

### Extraction of SNPs associated regulatory network in given tissue and two categories

Given a tissue, we check every RE in its regulatory network, if there was at least one SNP that was located in this RE, we extracted this RE and its upstream TFs and downstream TGs. We linked the SNPs in RE and this RE with edges and the SNPs, REs, TFs, TGs formed a SNP-associated regulatory network in a given tissue.

For 11 tissues of immune cells, we union the SNPs, REs, TFs, TGs. We assigned an edge to SNP-RE if it existed in at least one tissue’s SNP associated regulatory network, assigned an edge to TF-RE if it existed in at least one tissue’s SNP associated regulatory network, assigned an edge to RE-TG if it existed in at least one tissue’s SNP associated regulatory network. After the union and edge assignment, we constructed a regulatory network of immune cells. The regulatory network of epithelium cells was constructed with the same procedure.

## Data availability

The GWAS summary statistics of COVID-19 severity (“A2_ALL” study) was download at https://www.covid19hg.org/results/r4/. The GWAS summary statistics from WGS data were download at https://genomicc.org/data/. The constructed regulatory network atlas was freely available at https://github.com/AMSSwanglab. The eQTL dataset was download at the GTEx portal https://www.gtexportal.org/home/datasets. The collected paired expression and chromatin accessibility data was summarized in Table S1 and Table S2.

## Acknowledgements

We acknowledge funding from the Strategic Priority Research Program of the Chinese Academy of Sciences (XDPB17), National Key Research and Development Program of China (2020YFA0712402), and the National Natural Science Foundation of China (grants 12025107, 11871463, 61621003, 11688101). The Genotype-Tissue Expression (GTEx) Project was supported by the Common Fund of the Office of the Director of the National Institutes of Health, and by NCI, NHGRI, NHLBI, NIDA, NIMH, and NINDS. The data used for the analyses described in this manuscript were obtained fromthe GTEx Portal under dbGaP accession number phs000424.v8.p2 on 12/09/2020.

